# Serine auxotrophy is a targetable vulnerability driven by PSAT1 suppression in AML

**DOI:** 10.1101/2025.05.13.651470

**Authors:** Ilias Sinanidis, Panagiotis Tsakiroglou, Benjamin Dubner, Rebecca Foertsch, Julia Gondek, InYoung Choi, Bogdan Paun, Liang Zhao, Gabriel Ghiaur, W. Brian Dalton

**Author notes:** These authors contributed equally (Ilias Sinanidis, Panagiotis Tsakiroglou). Correspondence W. Brian Dalton: The Sidney Kimmel Comprehensive Cancer Center at Johns Hopkins, 1650 Orleans St, CRBI Room 285, Baltimore, MD 21231; Ilias Sinanidis: The Sidney Kimmel Comprehensive Cancer Center at Johns Hopkins, 1650 Orleans St, CRBI Room 216, Baltimore, MD 21231.

## Abstract

Serine metabolism is of growing biologic and therapeutic interest in cancer. Upregulation of the serine synthesis pathway (SSP) can fuel tumor growth, and cancers with this phenotype are often sensitive to SSP inhibitors. In parallel, dietary restriction of serine and glycine (SG) can suppress some cancers, but the determinants of sensitivity to this approach are poorly understood. This is especially true in acute myeloid leukemia (AML), where serine metabolism has been less explored. We report that a subset of human AML cell lines and primary samples are completely dependent on external serine, known as serine auxotrophy. These leukemias consistently suppressed the SSP enzyme PSAT1, failed to synthesize serine, responded to SG restriction *in vivo*, and were rescued by restoring PSAT1. We also found that AML with an SF3B1 K700E mutation showed additional dependence on the SSP enzyme PHGDH, that SG restriction synergized with venetoclax in serine auxotrophic AML, and that MECOM rearrangement was strongly associated with PSAT1 suppression and serine auxotrophy. These findings define a metabolically distinct AML subtype and nominate it for targeting by SG restriction.

## Introduction

Research in the last two decades has revealed diverse metabolic reprogramming in cancer. Alterations of metabolism are determined by tissues of origin, mutations, acquired treatment resistance, or other properties, and some changes create targetable metabolic vulnerabilities (1–3). In AML, deciphering the metabolic reprogramming induced by IDH1/2 mutations led to pathophysiologic insights and effective inhibitors (4–7). However, much of the landscape of metabolic reprogramming in cancer remains unknown—including in AML (8,9). This is especially true for the serine synthesis pathway (SSP), of significant interest in cancer metabolism but not well characterized in AML (9,10).

The SSP consists of three enzymes that make serine: PHGDH, PSAT1, and PSPH (10). SSP upregulation is seen in several solid malignancies, with mechanisms linked to genetic events in some cases. These include PHGDH amplification (11,12); transcriptional upregulation of PHGDH and/or PSAT1 by mutations in KRAS (13), BRAF (3), APC (14), NRF2 (15), and TP53 (16,17); and decreased PHGDH protein degradation by PARKIN mutations (18). In AML, FLT3 mutations have been implicated in upregulating the SSP (19). SSP upregulation supports cancer proliferation by fueling anabolism, especially in tissues with low environmental serine, and it may provide a crucial source of alpha-ketoglutarate (10,12,20). Accordingly, forced SSP upregulation can promote—and its downregulation can inhibit—growth of certain cancers (2,20,21). In AML, PHGDH knockdown and knockout were reported to decrease the fitness of FLT3-ITD cell lines (19,22).

In parallel, low or absent serine synthesis has occasionally been observed in some cancer cell lines (12,23–25). Recently, Choi et al systematically showed that ER/PR+ breast cancer lacks serine synthesis due to silencing of PSAT1, a state likely inherited from the cell-of-origin luminal breast epithelium (2). Moreover, we and others found that mutations in the spliceosome gene SF3B1 caused RNA missplicing-induced downregulation of PHGDH and decreased serine synthesis in human isogenic cell models (26–28). However, for most cancers, including AML, the extent, consequences, and mechanisms of downregulated serine synthesis are unknown.

Interest in the SSP is also therapeutic. Some cancers with elevated SSP show sensitivity to PHGDH inhibitors (19–21,25,29). Conversely, dietary deprivation of serine (and glycine, as the latter produces the former *in vivo*) lowers circulating serine in mice, is well tolerated, and slows growth of some solid tumors (2,14,20,21,30,31). In ER/PR+ breast cancer, serine and glycine deprivation (-SG) was effective because of absent SSP activity, as its restoration rescued -SG growth—thereby demonstrating serine auxotrophy for this cancer (2,20). Notably, a recent clinical trial of a reduced SG diet during solid tumor chemotherapy showed decreased circulating serine levels and tolerability in cancer patients (32). Given this translational potential and the limited knowledge of downregulated serine synthesis in AML, we sought here to determine the landscape of dependence on external serine, SSP activity, and efficacy of serine deprivation in human AML.

## Methods

### Cell culture

Master cultures of leukemia cell lines used RPMI with 1% P/S and 20% FBS. GMCSF was added as follows: 5 ng/mL for AML193, 10 ng/mL for MUTZ3 and KASUMI3, and 20 ng/mL for YCUAML1 and OCIAML20 (the latter two were also grown on OP9 stromal cells). Primary AML cells were grown on OP9 stromal cells in RPMI with 20 ng/mL of TPO, GM-CSF, IL3, IL6, FLT3, and SCF, 500nM UM729, and 750nM StemRegenin1.

### Serine starvation

Viable cell number and percentage were assessed using a Vi-Cell Blue, which uses trypan blue exclusion. Change in cell growth in -S versus +S media was calculated as K₀ / Kₛ, where K₀ and Kₛ are the growth constants in -S and +S, respectively. OP9 were used only in YCUAML1 in a concentration of 3000 cells/well.

### Mouse Xenograft Studies

Leukemia cell line xenografts were inoculated via tail vein with 10^7^ cells (unless otherwise specified) in 100 μL of HBSS, while MOLM13 (scrambled and PSAT1 KO) and P31FUJ (with K200A or WT PSAT1 overexpression) xenografts were inoculated with 10⁶ cells. Mice were started on –SG or +SG pellets, as described (26), on the day of inoculation (unless otherwise specified) and continuing until death or euthanasia. For - SG diet plus venetoclax, the diet began on day 3, with venetoclax given at 20 mg/kg on day 3, 50 mg/kg on day 4, and 100 mg/kg on days 5–7, 10, and 11.

### Methylation-specific PCR (MSP)

Genomic DNA was isolated and bisulfite converted. “Unmethylation” primers were manually designed while “methylation” primers were used as described by Choi et al (2), (suppl. Table 1). Percent methylation was calculated as follows: %_meth_ = 100 / (1 + 2^meth-unmeth^), where *meth* and *unmeth* are Ct values from the respective primer pairs.

### Gene overexpression

10^6^ cells were plated in 24-wells with lentivirus and LentiBoost (1:100), and centrifuged at 1250 g for 90 mins at 37°C. The multiplicity of infection (MOI) for lentivirus carrying either WT or mut sequence of PSAT1 and PHGDH was 10. At 72h post-infection, cells were treated with G418 at 250 μg/mL and blasticidin S HCI at 3 μg/mL for PSAT1 and PHGDH, respectively. Selected cells were collected and confirmed for stable expression of PSAT1 and PHGDH with western blot.

### Gene knockout (KO)

Lentiviral transduction was as above. MOI for lentivirus carrying either gRNA_exon1, gRNA30 and scramble was 0.008 (suppl. Table 1). At 72h post-infection, cells were treated with puromycin at 1 μg/mL, and single-cell-derived clones were grown in 96-wells. Knockout clones were confirmed by PSAT1 western blotting and gDNA Sanger sequencing of inactivating indels in the PSAT1 gene.

**Additional methods can be found in the supplementary materials.**

## Results

### A subset of AML cell lines is dependent on extracellular serine

To characterize extracellular serine dependence in AML, we first assessed the growth of 29 human AML cell lines and 2 populations of normal bone marrow (NBM) CD34+ cells *in vitro* using RPMI with or without serine (+S or -S). In -S, 66% (19/29) of AML lines, as well as NBMs, showed modestly decreased proliferation, compared to +S [growth rate (GR) -S/+S = 0.71 and 0.66, respectively] (Figure 1A). We did not observe significant cell death in -S for these cells (Figure 1B). This pattern resembles SSP-competent cells of other lineages, where slower -S growth reflects NAD+ depletion and reduced purine synthesis from compensatory serine production (33). In contrast, 34% (10/29) of AML lines exhibited growth inhibition and cell death in -S (GR -S/+S = 0.02) (Figures 1A-B). Death was caspase-dependent, as it was mitigated by the pan-caspase inhibitor QVD (Figures S1A-B). Because functional consequences have been ascribed to differences between traditional cell culture media and formulations better approximating the human blood metabolome (29,34), we confirmed these results in human plasma-like media (HPLM), which produced GR -S/+S values of 0.56 and 0.05 for cells that had been independent or dependent, respectively, on external serine in RPMI (Figure S2A). These data indicate that AML cell lines show a heterogeneous response to serine restriction, and there is a distinct subset that exhibits serine auxotrophy.

**Figure 1.**
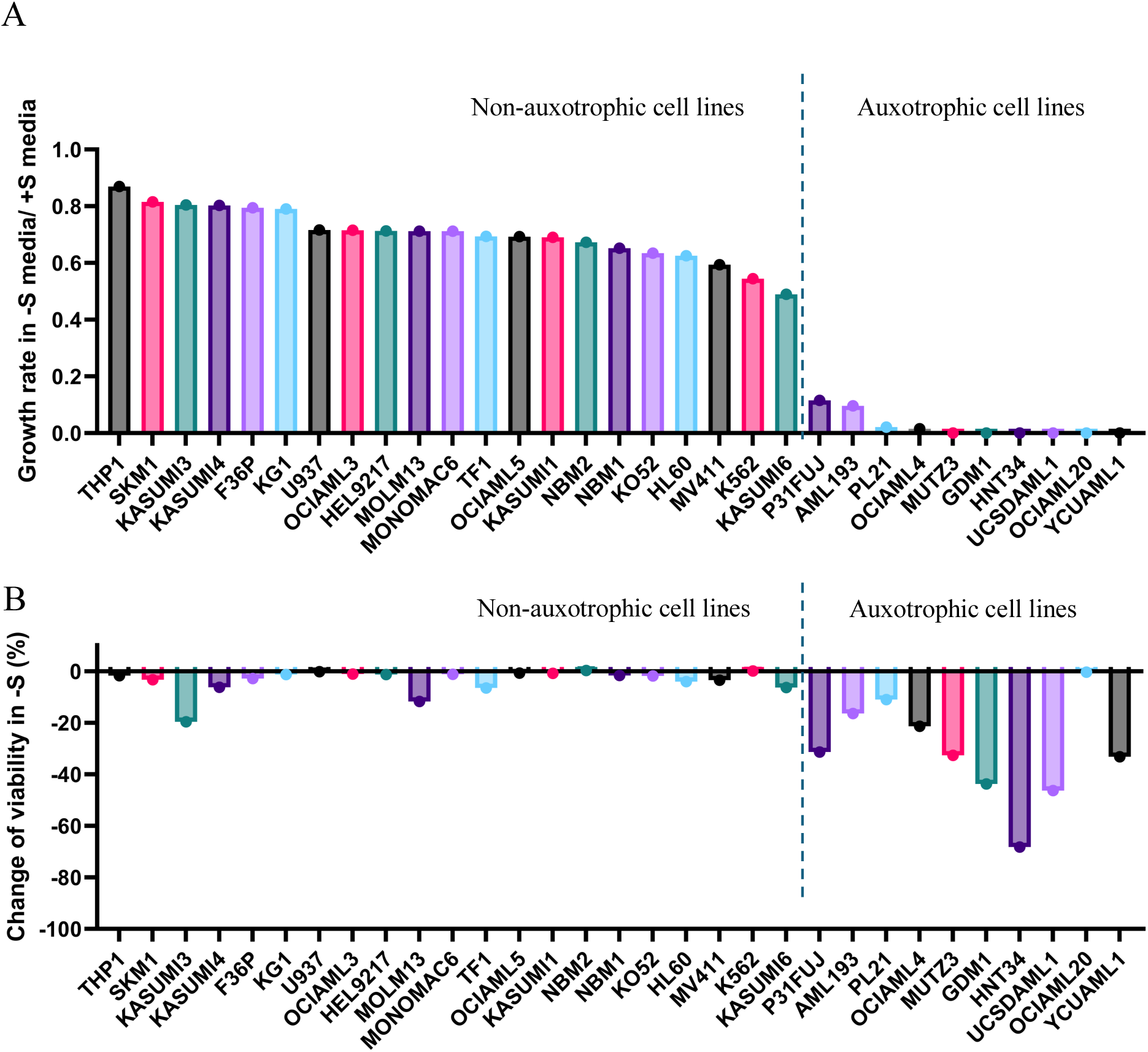
Landscape of dependence on external serine in AML cell lines. (A) The ratio of growth rate in -S versus +S RPMI was determined by triplicate viable cell counts in 3 timepoints over at least 2 population doublings in +S media. (B) Percent change of viability (V) calculated using trypan blue exclusion. %V_change_ = 100% x (V_-S_ – V_+S_)/V_+S_, where V_-S_ and V_+S_ are the viability at the last timepoint of Figure 1A starvation curves in –S and +S RPMI, respectively.

### PSAT1 is suppressed in serine auxotrophic AML cell lines

To investigate mechanisms of serine auxotrophy, we compared gene expression profiles of functionally defined serine auxotrophic (SA) and non-auxotrophic (NA) cell line groups using RNA-seq from the Cancer Cell Line Encyclopedia (CCLE). This revealed that two outlying genes were profoundly downregulated in SA cells: MGST1 and PSAT1 (log2 SA/NA = −4.9 and −5.1, respectively) (Figure 2A). PSAT1 is the second enzyme in the SSP (10), so its downregulation could compromise serine synthesis and induce serine auxotrophy (Figure 2B). We therefore measured PSAT1 protein with western blotting, which showed variable but present expression in all 19 NA cell lines (Figures 2C, and S2B). Conversely, no PSAT1 protein was detected in any conditions in 9 out of 10 SA lines, with a trace amount seen in the 10^th^ (OCIAML20) (Figures 2C, and S2C). By contrast, the protein level of PHGDH, the first SSP enzyme, was variably detected by western blot and did not strongly correlate with dependence on external serine, as did PSAT1 (Figure 2B). Thus, PSAT1 expression is conspicuously absent (or very low) in SA AML cell lines.

**Figure 2.**
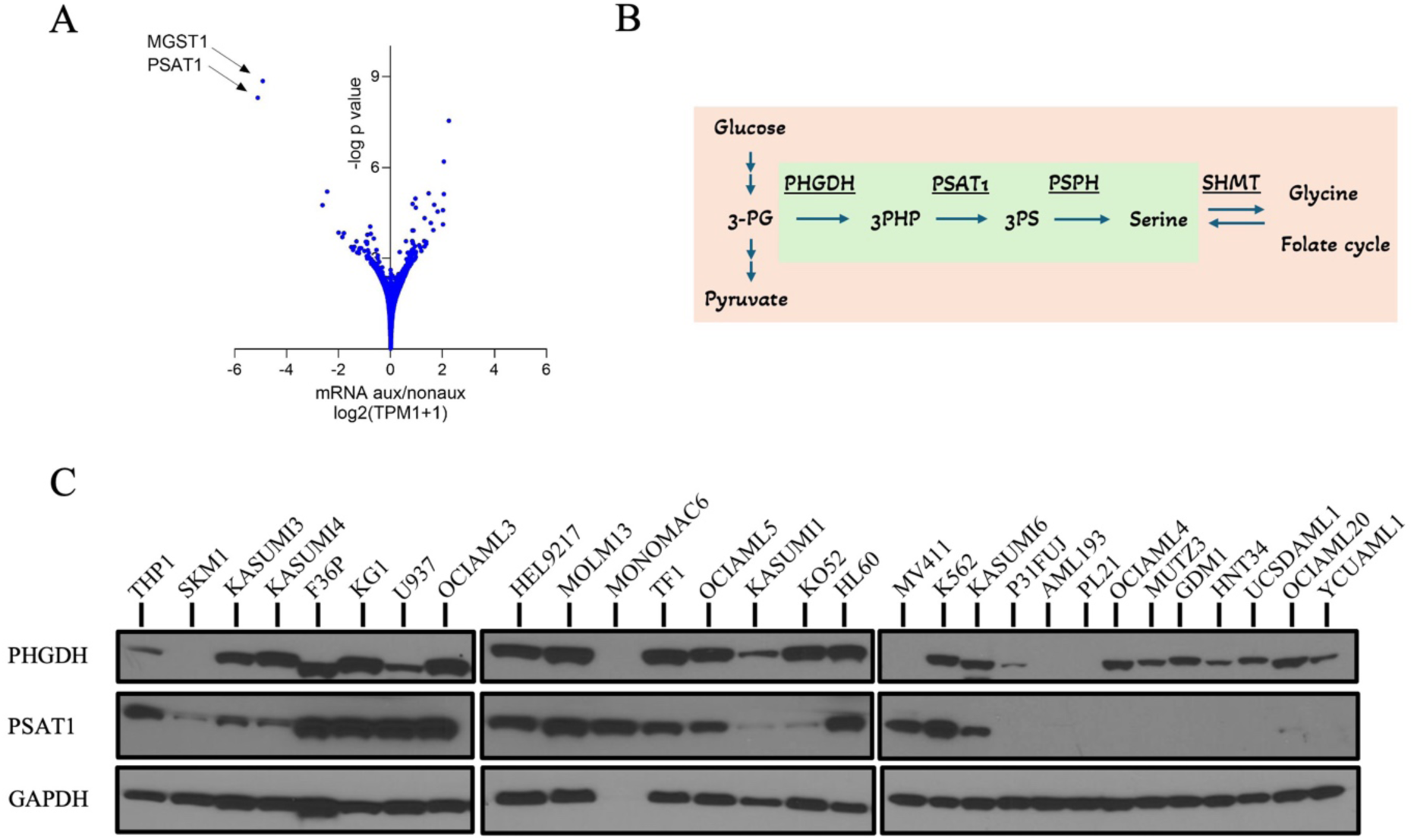
PSAT1 is suppressed in serine auxotrophic AML cell lines. (A) Gene expression profiles of functionally defined auxotrophic and non-auxotrophic AML cell lines, analyzed using RNA-seq data from the CCLE. (B) Schematic representation of the serine synthesis pathway (SSP) highlighted in green. The SSP connects glycolysis with one-carbon metabolism and glycine synthesis. (C) PSAT1 and PHGDH protein levels in AML cell lines. GAPDH serves as the loading control. MONOMAC6 expression was validated separately using COX-IV (Figure S2B).

### Serine auxotrophic AML cell lines do not synthesize serine

To determine serine synthesis capacity of SA cell lines, we fed cells ^13^C-glucose and quantified synthesized M+3 serine by mass spectrometry in +S and -S conditions. The NA cell lines MOLM13 and OCIAML3 exhibited low baseline serine synthesis (0.8% and 4.6%, respectively) in +S, consistent with SSP-competent cells from other lineages, which primarily use imported serine when it is available (Figure 3A) (12,33,35). After 6h of -S, MOLM13 and OCIAML3 increased M+3 serine to 12.5% and 8.2%, respectively, indicating an intact capacity to sense and respond to serine depletion (Figure 3A). Conversely, the SA cell lines P31FUJ, UCSDAML1, YCUAML1, and MUTZ3 synthesized no detectable serine at baseline or in -S (Figure 3A). Their sensing of serine depletion appeared operational, as the metabolic stress-induced transcription factor ATF4 was upregulated in -S (Figure 3B). However, PSAT1 was not induced, a known consequence of ATF4 upregulation (Figure 3B) (15,36,37). These data demonstrate that SA cell lines do not synthesize serine to compensate for its loss from the environment.

**Figure 3.**
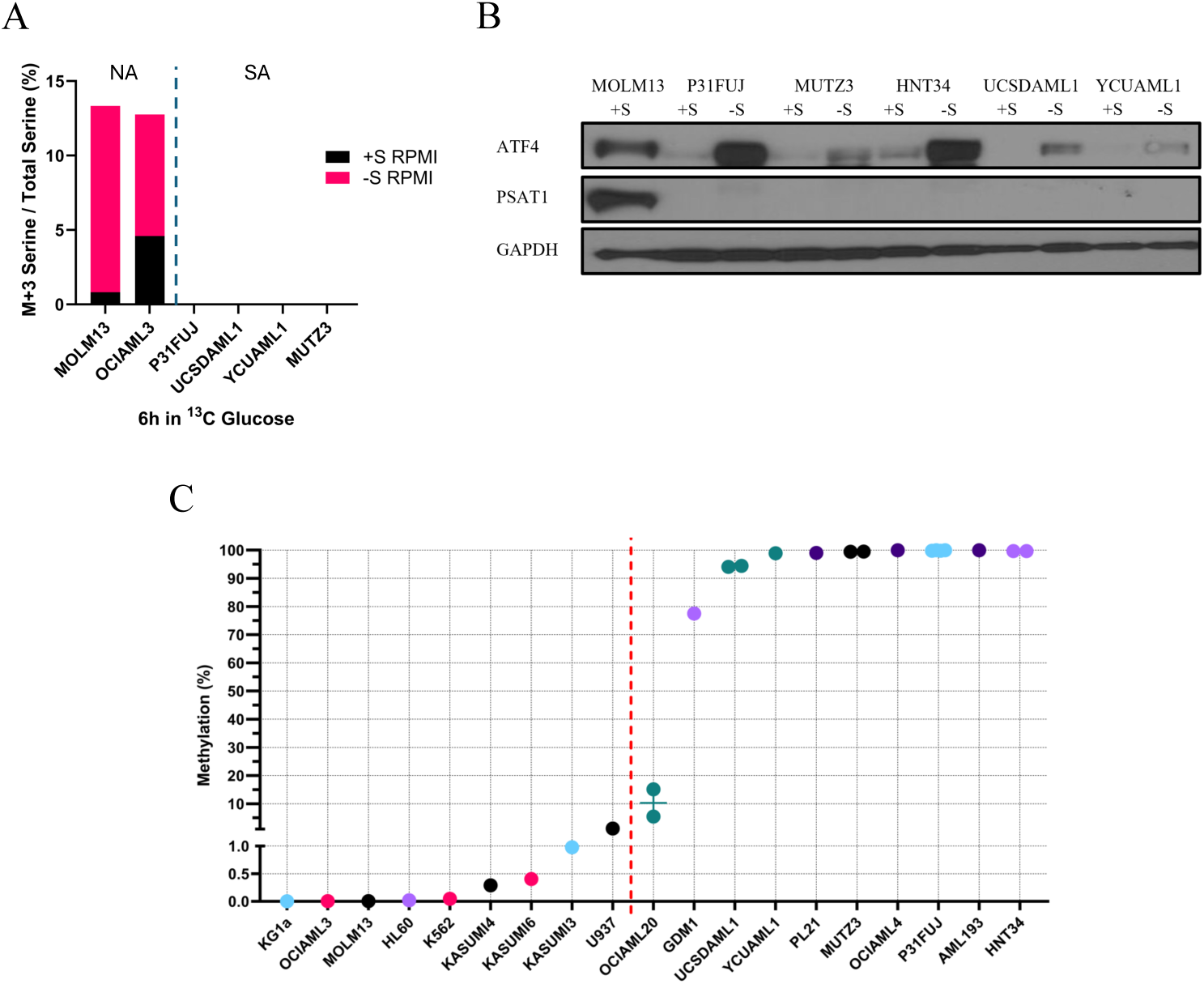
Serine auxotrophic AML cell lines sense –S metabolic stress but do not synthesize serine. (A) De novo M+3 serine synthesis using ¹³C labeling after a 6-hour incubation in ¹³C-glucose media. NA: non-auxotrophic cell lines. SA: auxotrophic cell lines. Values represent the mean of quadruplicates. (B) Western blot showing ATF4 and PSAT1 expression levels after 24 hours of incubation in −S or +S RPMI. GAPDH serves as the loading control and MOLM13 sample as a PSAT1 positive control. (C) Baseline methylation percentage (%) of AML cell lines, calculated based on Ct values of methylation -specific and unmethylation-specific qPCR amplification.; %meth = 100 / (1 + 2^meth-unmeth^). Each sample was analyzed in triplicate.

### Serine auxotrophy is associated with PSAT1 promoter methylation

We next interrogated the DNA methylation status of the PSAT1 promoter, as this was found to mediate silencing of this gene in ER/PR+ breast cancer (2). Using quantitative MSP, we found that NA AML cell lines had absent or very low PSAT1 promoter methylation (Figure 3C). In contrast, 9 out of 10 SA cell lines showed high (75-100%) levels of methylation (Figure 3C). The 10^th^ SA cell line had ∼10% methylation, which, while higher than all NA cells, was lower than other SA cells (Figure 3C). Of note, this cell line, OCIAML20, is also the auxotrophic cell line with very low, but still detectable, PSAT1 protein, potentially consistent with partial gene silencing (Figure S2C). These findings suggest that promoter methylation may contribute to the suppression of PSAT1 in SA AML cell lines.

### PSAT1 determines serine auxotrophy in AML cell lines

We next tested the functional role of PSAT1 in SA for AML cell lines. PSAT1 knockout with two different sgRNAs in the NA cell line MOLM13 rendered cells SA, confirming PSAT1 is indeed necessary for -S growth in this context (Figures 4A, and S3A). To determine whether PSAT1 silencing is the main mediator of native SA in AML cell lines, we re-expressed PSAT1 via lentiviral cDNA delivery in 5 auxotrophic AML lines. Enzyme-dead but structurally intact K200A PSAT1 was used as a control due to nonenzymatic functions of the protein (38,39). In 3 of these lines (UCSDAML1, P31FUJ, MUTZ3), re-expression of WT, but not K200A, PSAT1 was sufficient to rescue SA (GR -S/+S = 0.93 and 0.15 for WT and K200A, respectively) (Figures 4B, and S3B). In PL21 and HNT34 cells, PSAT1 re-expression provided a more partial rescue of auxotrophy (GR -S/+S = 0.61 and 0 for WT and K200A, respectively) (Figures 4C-D, and S3B-C). HNT34 harbors a K700E mutation in SF3B1, which we previously found caused RNA missplicing-induced downregulation of PHGDH and lower serine synthesis in isogenic human cell models (26). We therefore also overexpressed PHGDH in HNT34 cells to test its role in serine auxotrophy. Enzyme-dead D175N/R236K/H283A PHGDH was used as a control due to noncatalytic functions of this protein (40–43). Interestingly, HNT34 cells with double transduction of WT PSAT1 and PHGDH exhibited full rescue of auxotrophy, while overexpression of WT PHGDH with K200A PSAT1 had no impact (Figures 4D). Moreover, isotopomer tracing of ^13^C-glucose showed no M+3 serine in HNT34 cells expressing K200A PSAT1, regardless of PHGDH status. Expression of WT PSAT1 and enzyme-dead PHGDH induced some SSP activity (M+3 = 0.48%), but serine synthesis was further increased 4.6-fold in cells with double rescue of WT PSAT1 and PHGDH (M+3 = 2.25%) (Figure 4E). This points to a deficiency in both the first and second steps of the SSP in SF3B1^K700E^ HNT34 cells. In contrast, WT PHGDH overexpression had no additional effect on -S growth in P31FUJ and UCSDAML1 (Figures S3B, and S4A-B). We also re-expressed MGST1, a glutathione transferase implicated in ferroptosis (44) that was the other highly downregulated gene in our transcriptome analysis (Figure 2A), in HNT34 and MUTZ3 cells, but this did not affect auxotrophy (data not shown). These findings indicate that PSAT1 suppression is a key mechanism of SA in AML cell lines—and that additional factors, such as missplicing-induced PHGDH downregulation by SF3B1^K700E^, can also contribute to the phenotype.

**Figure 4.**
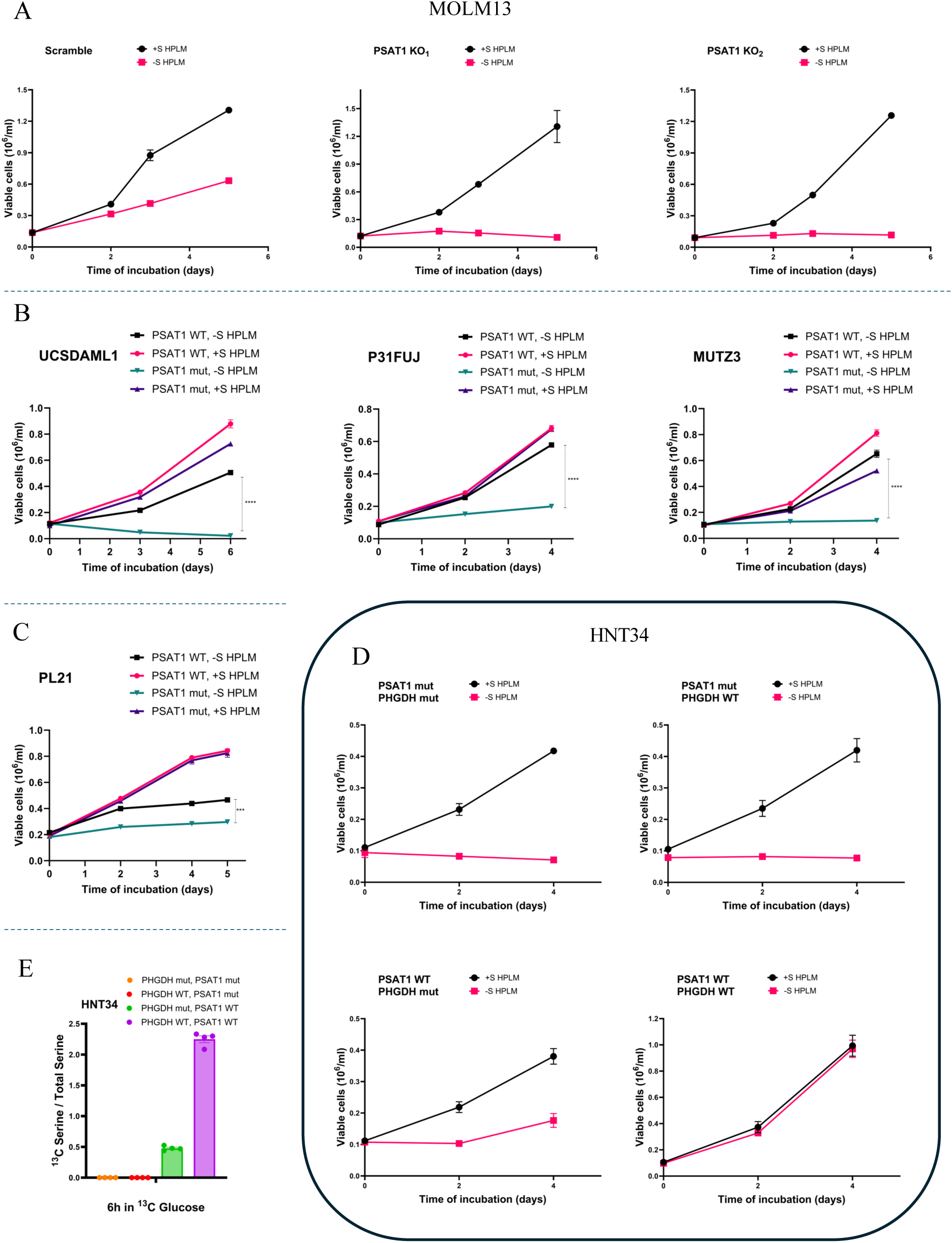
PSAT1 determines serine auxotrophy in AML cell lines. Serine starvation curves of (A) MOLM13 PSAT1 knockouts, (B-C) UCSDAML1, P31FUJ, MUTZ3, and PL21 with PSAT1 overexpression in +S or -S HPLM media. Each sample was run in triplicate. (D) Serine starvation curves of HNT34 cell lines restoring the expression of PSAT1 and/or PHGDH in +S or -S HPLM. Each sample was assessed in quadruplicate. (E) De novo serine synthesis in HNT34 with overexpression of PSAT1 and/or PHGDH using ¹³C labeling after a 6-hour incubation in ¹³C-glucose media. Values represent the mean ± SEM of quadruplicates. (B-E) “Mut” refers to inactivation mutations of the PSAT1 or PHGDH protein. ***p<0.001, ****p<0.0001.

### Dietary SG restriction is therapeutic in auxotrophic AML cell lines

Dietary SG restriction (-SG diet) decreases these amino acids by 40-60% in mice (and more recently humans), is nontoxic, and inhibits certain solid tumors, but its activity in AML is unknown (32,35,45). We tested whether -SG diet exerts antileukemic activity against auxotrophic AML lines engrafted in NSG mice. We found that auxotrophic P31FUJ cells produced a highly proliferative leukemia that was fatal within 2-3 weeks, and -SG diet significantly extended survival and decreased leukemic burden of engrafted mice (Figures 5A-B). The auxotrophic cell lines YCUAML1, PL21, and OCIAML20 produced leukemias that were fatal in 8, 6, and 12 weeks, respectively, and -SG diet extended survival by ∼2 weeks for each of these (Figure 5A). These results indicated that SA leukemias are targetable and can be inhibited by dietary SG restriction.

**Figure 5.**
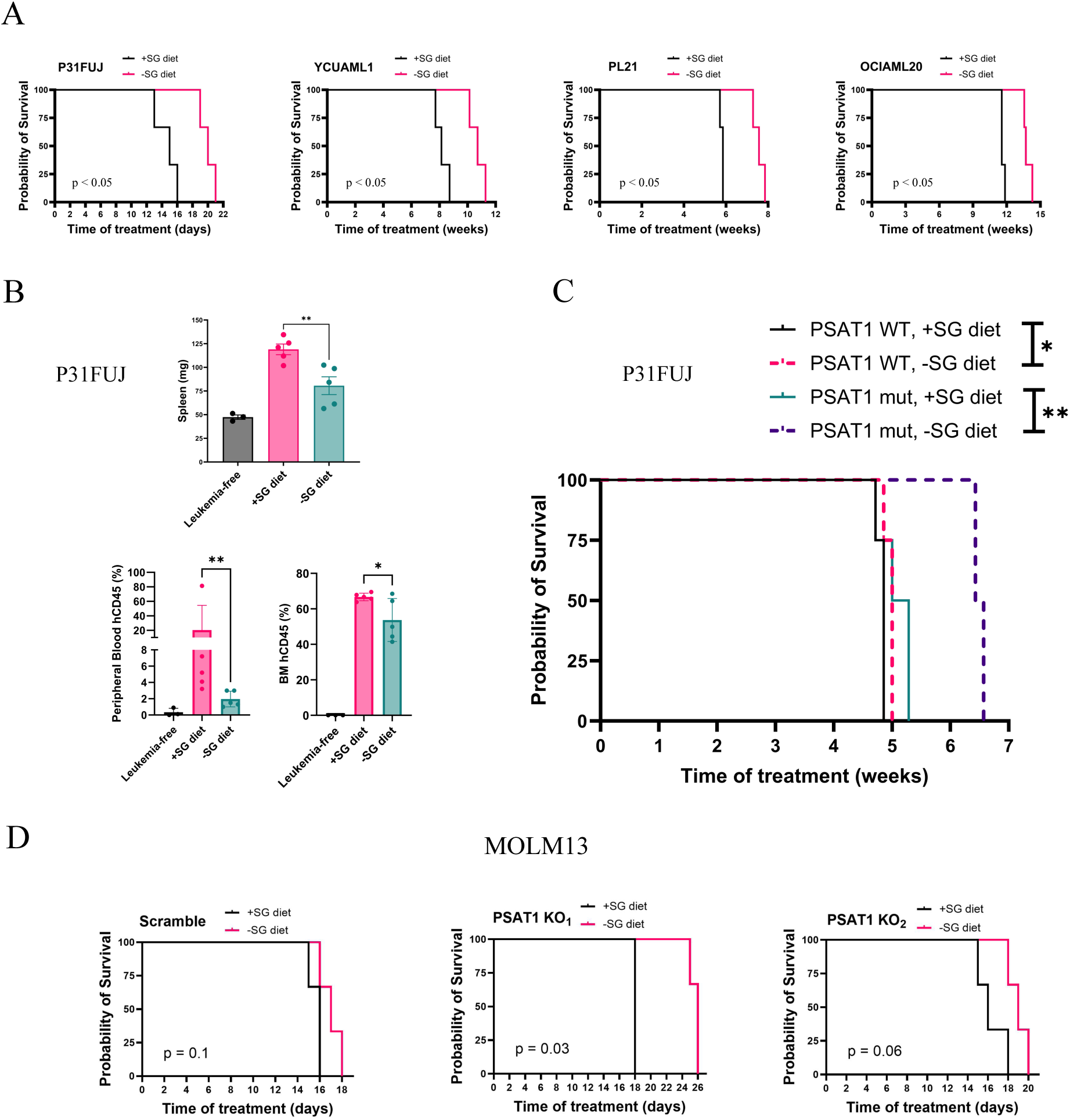
Serine and Glycine-restricted (-SG) diet extends survival and decreases leukemia burden in AML xenografts. (A) Kaplan–Meier survival curves of AML xenografts inoculated with 10^7^ cells per mouse (n = 3 per group). (B) Leukemia burden, assessed through spleen weight, peripheral blood human hCD45% levels, and bone marrow hCD45%, in P31FUJ xenografts. Mice were euthanized 14 days after cell inoculation. N = 5 per study group, n = 3 for leukemia free control group. Values represent means ± SEM. (C) Kaplan-Meier survival curves of xenografts bearing P31FUJ leukemia expressing WT or K200A (mut) PSAT1. 10^6^ cells were inoculated per mouse (n = 4 per group). (D) Kaplan–Meier survival curves of MOLM13 xenografts inoculated with 10⁶ cells per mouse (n = 3 per group). (A-D) Treatment with -SG or +SG diet was initiated on the day of inoculation and continued until mouse death or euthanasia. *p<0.05, **p<0.01

To evaluate the role of PSAT1 in the response to -SG diet, we engrafted mice with our P31FUJ cells modified to express WT or K200A PSAT1. Mice bearing K200A-expressing P31FUJ cells, like parental cells, showed increased survival with -SG (Figure 5C). Interestingly, WT PSAT1-expressing P31FUJ mice lost sensitivity to -SG, surviving only ∼1 day longer than +SG (Figure 5C). Similarly, survival of mice engrafted with NA (PSAT1 expressing) MOLM13 cells was not significantly affected by -SG diet (Figure 5D), while auxotrophic PSAT1-KO MOLM13 xenografts exhibited improved survival with -SG (Figure 5D). These data show that PSAT1 expression is a determinant of sensitivity to dietary SG restriction in AML.

### Venetoclax synergizes with serine deprivation

The BCL2 inhibitor venetoclax (VEN), together with a hypomethylating agent or low-dose cytarabine, is the standard of care for older or unfit AML patients, and it is being evaluated as a “universal sensitizer” in many combinations in clinical trials (46). We therefore investigated combining serine restriction with VEN in auxotrophic AML, as well as the role of PSAT1 in mediating any effects. In +S, both P31FUJ and PL21 cells (expressing K200A PSAT1) exhibited some sensitivity to 30-100nM VEN, but cell death increased an additional ∼50% in -S with VEN (Figure 6A), indicating synergy between -S and VEN in SA cells. Re-expression of WT PSAT1 partially rescued viability of cells treated with -S and VEN (Figure 6A), pointing to a lack of serine synthesis as a driver of this synergy. To test activity *in vivo*, we treated P31FUJ-engrafted mice with +/-SG diet and +/-VEN. Three of four mice that received -SG diet + VEN lived the longest of all mice, although the fourth died soon after treatment initiation (Figure 6B). This suggests both efficacy and toxicity of -SG diet + VEN, and the early death pattern might be potentially consistent with tumor lysis syndrome, a known consequence of VEN in humans that has been seen with another effective VEN combination in mice (47). Overall, these findings point to synergy between serine deprivation and VEN in SA AML, but they also emphasize the need to consider dosing optimization to mitigate toxicity.

**Figure 6.**
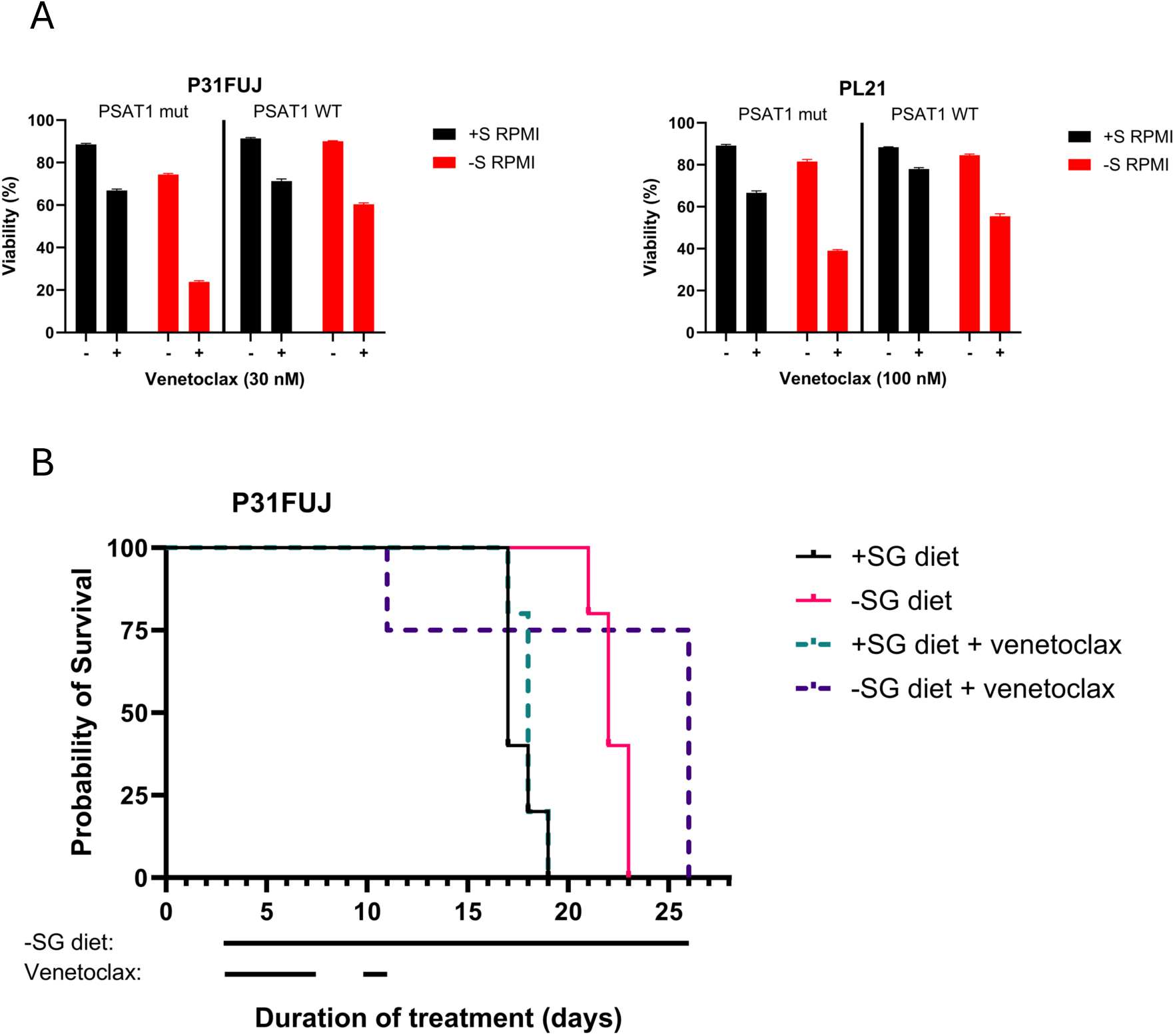
Combination treatment with –SG and venetoclax. (A) P31FUJ and PL21 cell viability after 24h and 48h in –S RPMI and/or venetoclax, respectively. Values represent means ± SEM. (B) Kaplan–Meier survival curves of P31FUJ xenografts inoculated with 10^7^ cells per mouse (*n* = 5 per group, n = 4 in combination therapy arm). The testing diet was initiated on day 3 and continued until mouse death or euthanasia. Venetoclax was administered on day 3 at a dose of 20 mg/kg, on day 4 at a dose of 50 mg/kg, and on Days 5, 6, 7, 10, and 11 at 100 mg/kg.

### PSAT1 expression in genetic subtypes of AML

We next focused on how our cell line findings relate to primary human AML samples. As diagnosis, prognosis, and treatment of AML are all strongly guided by its genetic drivers, we analyzed the landscape of PSAT1 expression across different AML mutations and cytogenetic aberrations using RNA-seq from TCGA, BEAT-AML, and Leucegene patient cohorts. PSAT1 downregulation was associated with MECOM rearrangement (i.e. inv3), CBFB-MYH11 fusion (i.e. inv16), and mutations in GATA2, WT1, and CEBPA in two or more cohorts (Figures 7A-C). The association with MECOM rearrangement (MECOMr) was especially notable, as MECOMr AML is a highly treatment-refractory and deadly AML. Moreover, 71% (5/7) of MECOMr cell lines exhibited PSAT1 silencing and SA (Figure 7D), suggesting sensitivity to serine restriction may be particularly high for this most lethal AML. In contrast, PSAT1 upregulation was associated with mutations in TP53 and U2AF1 in two or more patient cohorts (Figures 7E, and S5A). Additional associations in BEAT-AML included slightly lower PSAT1 with mutations in NPM1 and FLT3 and higher PSAT1 with KMT2A rearrangement and MPL mutation (Figure S5A). DNA methylation data were available from TCGA and BEAT-AML, and PSAT1 methylation was elevated in WT1- and CEBPA-mutated AML, and, in BEAT-AML, inversely correlated with TP53 mutation (Figures 7A, 7C, 7E and S5B). These data show that PSAT1 expression is heterogeneous yet consistently correlated with distinct genetic drivers in primary AML samples.

**Figure 7.**
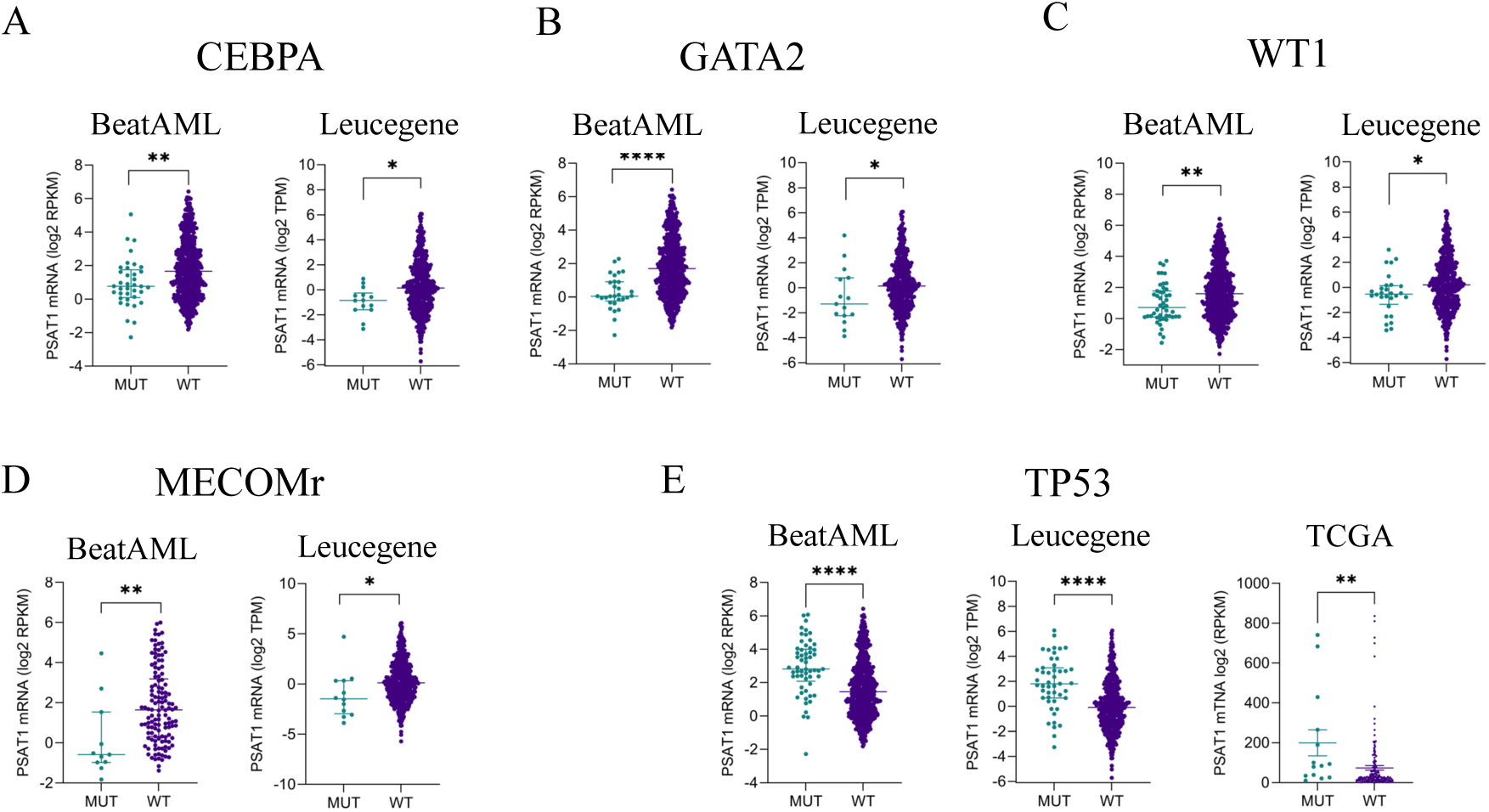
PSAT1 mRNA expression in AML patient cohorts. (A) CEBPA-, (B) GATA2-, (C) WT1-, (D) MECOMr-, and (E) TP53-mutated AML samples from RNA-seq in Leucegene, BEAT-AML, and/or TCGA patient cohorts. Panels (A-E) show median values with interquartile ranges. *p<0.05, **p<0.01, ***p<0.001, ****p<0.0001

### Primary AML samples with silenced PSAT1 are sensitive to serine deprivation

Finally, we functionally assessed *in vitro* external serine dependence and PSAT1 expression in nine primary human AML samples (Figure 8A). In an effort to capture the full spectrum of PSAT1 expression, we included samples bearing genetic aberrations that correlated with differing levels of PSAT1 in our transcriptional analysis above. Protein levels of PSAT1 and growth in -S media indeed fell along a spectrum and largely correlated (Figures 8A-C). At one end, three samples showed PSAT1 levels higher than NBM and robust growth in -S media (Figures 8A-B). At the other, two samples showed undetectable PSAT1 and complete serine auxotrophy (Figures 8A-B, and S6A). Three other patients had very low PSAT1 and slight or no growth in -S media (Figures 8A-B). The last patient, p4319, was an exception, with high PSAT1 but only slight growth in -S media (Figures 8A-B). Of note, the two auxotrophic, PSAT1-undetectable samples both had MECOMr, and a NA, PSAT1-high sample had TP53 mutation, consistent with patient cohort data (Figure 7D-E, 8A and S6B). Together, these data demonstrate that, like AML cell lines, primary samples exhibit heterogeneous dependence on external serine, and there is a subset of primary AML with absent (or very low) PSAT1 and serine auxotrophy.

**Figure 8.**
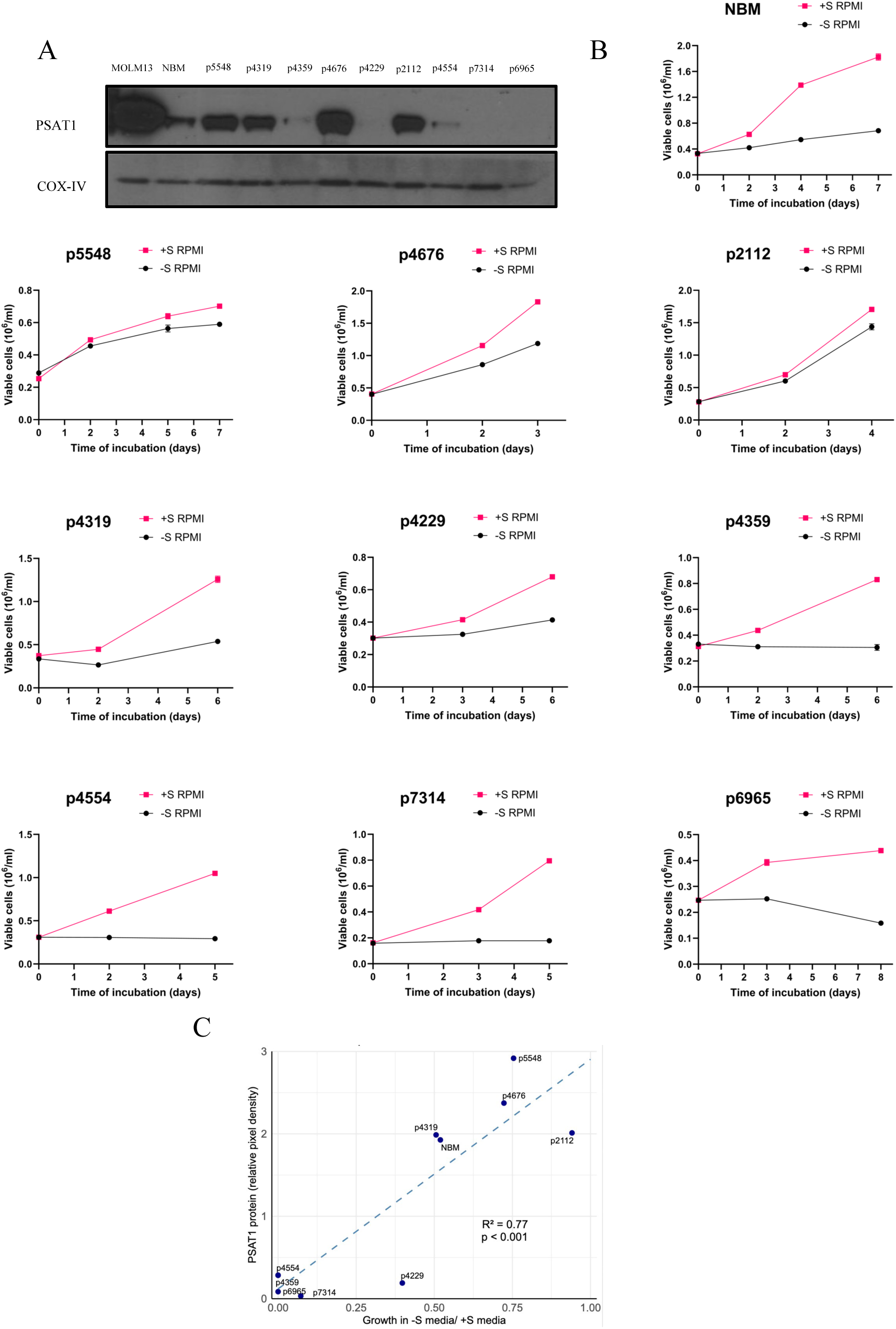
PSAT1 expression and dependence on external serine in primary AML samples. (A) Western blot showing the PSAT1 protein expression in primary AML patient samples. COX-IV serves as the loading control, and MOLM13 as the positive PSAT1 control. (B) Serine starvation curves of primary AML patient samples in +S or -S RPMI. Media was enriched with TPO, GM-CSF, IL-3, IL-6, FLT3-L, SCF, UM729, and SR-1. Values represent the mean ± SEM of triplicates. (C) Correlation plot between PSAT1 protein levels (measured by western blot densitometry) and ratio of cell growth rate, –S/+S, calculated from starvation curves. The data points of p4554 and p4359 overlap in the graph.

## Discussion

We report that a subset of AML exhibits serine auxotrophy mediated by silencing of the PSAT1 gene and targetable with dietary SG restriction (+/-venetoclax). These findings have implications for understanding and targeting altered serine metabolism in cancer. First, the presence of SA in AML expands the known landscape of this phenomenon, which has been characterized in ER/PR+ breast cancer and noted occasionally in other cell lines. As Sullivan et al found that serine-low, but not high, murine tissues encourage SSP upregulation in cancers arising from them, our results would suggest bone marrow is serine-rich enough to support outgrowth of auxotrophic cells (20). To what extent other tissues enable auxotrophic cancers is unclear, as is the question of whether SA is itself a cancer driver in any contexts or is simply a passenger effect. What is clear from our data, however, is that this subset of AML is highly reliant on exogenous serine—even for cell viability. To our knowledge, this has not been reported in other cancers, which undergo cell cycle arrest, rather than cell death, in similar culture conditions. Auxotrophic AML may therefore be especially, if not uniquely, dependent on external serine among human cancers.

Second, these results further credential the role of PSAT1 in determining external serine dependence in cancer. Canonically, PHGDH has been considered the rate-limiting step of the SSP, and apart from ER/PR+ breast cancer, most instances of SSP suppression in other cell lines involved PHGDH downregulation (12,23–25). However, like ER/PR+ breast cancer, SA in AML was remarkably and consistently associated with PSAT1 suppression, and its re-expression provided rescue of SA in multiple AML cell lines. We did find that the SF3B1^K700E^ cell line HNT34, which has RNA missplicing-induced downregulation of PHGDH, also required PHGDH overexpression for robust growth without exogenous serine. However, auxotrophy in the SF3B1^K666N^ cell line MUTZ3 was largely rescued by PSAT1 alone, indicating its downregulated PHGDH (Figure 4B) contributed little, if at all, to its auxotrophy. This difference may be due to the lower magnitude of RNA missplicing generally—and in PHGDH specifically—by the K666N vs. K700E mutations in SF3B1 (48–50). That said, two primary AML samples also had SF3B1 mutation, one PSAT1-negative and auxotrophic and the other PSAT1-positive and NA (Figure 8), further illustrating the primacy of PSAT1 in determining serine auxotrophy in AML.

As to the mechanisms responsible for PSAT1 downregulation in SA AML, our data point to DNA promoter methylation as a contributing factor. The inverse correlation between PSAT1 promoter methylation and protein expression was very strong in AML cell lines, though it was less pronounced in primary AML samples (Figure S6C). This may suggest there are additional epigenetic mechanisms of PSAT1 silencing and/or different patterns of CpG island methylation for PSAT1 in primary AML. Moreover, the underlying causes for PSAT1 methylation (or other epigenetic silencing mechanisms) are unclear at this time. While PSAT1 silencing may be a pre-existing state of the leukemia cell-of-origin, it may also be a more direct downstream consequence of the genetic drivers associated with SA AML (Figure S5B). Investigating mechanistic links between PSAT1 suppression and genetic aberrations like MECOMr, WT1 mutation, and others, is clearly an area for future work.

A third implication of our findings is for therapeutic serine deprivation strategies. Treatments with a -SG diet have been tested in solid tumor models (2,14,20,21,31,51–54). However, it has not been previously studied in leukemia, and we found that a -SG diet extended the survival of mice engrafted with four different auxotrophic, PSAT1-suppressed AML cell lines, as well as MOLM13 AML cells made auxotrophic through PSAT1 knockout. Together with our validation of serine auxotrophy in PSAT1-suppressed primary AML samples, these findings nominate AML as a cancer with substantial potential for therapeutic susceptibility to dietary serine restriction. Moreover, we found that venetoclax potentiated the efficacy of serine deprivation, albeit with a toxicity concern *in vivo*. Further studies will be necessary to determine whether dose optimization avoids this toxicity. Other therapy-related subjects for future work include studies in immunocompetent mice, as -SG can modulate antitumor immunity (32); other -SG and antileukemia therapy combinations; and mechanistic studies into the synergy of -S with venetoclax, which may involve reported direct effects of VEN on metabolism (55,56).

Finally, our results may directly inform potential clinical trials. Tong et al (32) recently reported results of a single-arm phase I study of 20 solid tumor patients undergoing chemotherapy who received a low-protein, fruits-and-vegetables diet supplemented with -SG nutritional powder. This successfully lowered serum SG by ∼50%, similar to mice, and produced no serious toxicities. Our work here supports the idea that testing this or other dietary approaches to restricting serine (for example, SG-depleted medical food, NCT05078775) is warranted in AML patients. Furthermore, our data nominate a potential biomarker, PSAT1, whose expression may have predictive value for treatment response and patient selection. While protein biomarkers are more complicated to integrate clinically than already-available genetics, there are several successful examples, such as PD1, PDL1, HER2, ER/PR, and others. Moreover, genetic aberrations associated with PSAT1 suppression could also potentially be incorporated into -SG studies to refine response prediction and patient selection. Particularly compelling was the association of TP53 mutations with elevated PSAT1 in all data sets—as well as high PSAT1 and robust growth in -S of our TP53-mutant primary sample (Figures 7E, 8A-C, and S5B). This would, unfortunately, argue for the exclusion of this poor prognosis AML in -SG dietary studies. However, the association of MECOMr with undetectable PSAT1 and complete SA was also striking—and would argue for specific inclusion of this poor prognosis AML in -SG studies. In conclusion, our study reports a hitherto uncharacterized metabolic vulnerability with translational potential for a subset of human patients with AML.

## Acknowledgments

We would like to thank Teodora Supeanu for her guidance on the animal experiments, and Ioannis Evangelos Louloudis for his assistance in generating the graph in Figure 8C using R. This research was supported by National Heart Lung and Blood Institute at the National Institutes of Health (R01HL159306), Leukemia and Lymphoma Society (6631–22), Edward P. Evans Foundation (138162), Maryland Stem Cell Research Fund (MSCRFL-5650), and Gabrielle’s Angel Foundation for Cancer Research (133).

## Authorship Contribution

Conceptualization and research design: W.B.D. and I.S, data collection and analysis: W.B.D and I.S, animal experiments: I.S., P.T., and B.D., funding acquisition: W.B.D., performed research: I.S., P.T., B.D., R.F., J.G., and I.C., patients’ samples collection, and processing: B.P. and G.G., metabolomic analysis: L.Z. and P.T., supervision: W.B.D., writing: original draft: I.S., W.B.D., P.T., and L.Z., and writing: review & editing: all authors.

