## Supplementary materials for "Serine auxotrophy is a targetable vulnerability driven by PSAT1 suppression in AML"

Supplementary Methods p2-5

Supplementary Tables p6-8

Supplementary Figures p9-15

References p16

### Supplementary Methods

**Cell lines and AML patient samples:** THP1, KASUMI3, KASUMI4, KASUMI6, AML193, TF1, KG1, U937, HEL9217, GDM1, and OP9 were purchased from ATCC. KASUMI1, HL60, K562, MOLM13, MONOMAC6, and MV411 were gifts from LP Gondek (Johns Hopkins University). SKM1, F36P, OCIAML3, OCIAML5, PL21, OCIAML4, MUTZ3, HNT34, and UCSDAML1 were purchased from DSMZ. KO52 was purchased from AcceGen. P31FUJ was purchased from the JCRB cell bank. OCIAML20 was a gift from M Minden (Princess Margaret Cancer Centre)(1). YCUAML1 was a gift from H Kunimoto (Yokohama City University) (2). STR authentication and mycoplasma testing were performed for all cell lines. Primary samples were collected from the bone marrow or blood of AML patients who provided written consent under the J0969 institutional review board–approved cell bank protocol at Johns Hopkins University. In all primary cultures, high blast number (>75%) was confirmed by a CD45-dim, low-side scatter phenotype by flow cytometry.

**Cell line transcriptomic analysis**. For auxotrophic and non-auxotrophic (as defined in Figure 1) cell lines that were also part of the Cancer Cell Line Encyclopedia, mRNA gene expression profiles of the two groups were subjected to a custom two-class comparison in the DepMap portal (https://depmap.org/portal/). A volcano plot of the difference in gene expression (auxotrophic log2(TPM1+1) – non-auxotrophic log2(TPM1+1)) versus -log10(p-value) was created using PRISM.

**Serine starvation studies:** For -S studies, we used two media: commercially available serine-, glycine-, and dextrose-free RPMI from Teknova, and customized serine- and glycine-free HPLM from ThermoFisher. For RPMI, 11 mM dextrose, 133 µM glycine, +/- 286 µM L-serine were added back to the media, along with 20% dialyzed FBS (dFBS) and 1% Pen/Strep. For HPLM, 300 µM glycine +/- 150µM L-serine were added back, along with 5% dFBS and 1% Pen/Strep. Growth factors were then added as in master cultures per above. Cells were seeded at $1 \times 10^5$ cells/mL for cell lines and $2–3 \times 10^5$ cells/mL for primary AML samples in flat-bottom 96-well plates, in triplicate or quadruplicate. To prevent cell death under -S conditions, $1 \times 10^6$ HNT34 cells were seeded into 6-well plates and incubated for 3 days in -S or +S media, with or without 25 µM pan-caspase inhibitor QVD. On day 3, the cells were resuspended in

Propidium Iodine (3 μg/ml) and run on an LSR-II flow cytometer. Exponential (Malthusian) growth constants (K) were calculated using PRISM (GraphPad).

**Western blotting:** Total protein was extracted from 1 × $10^6$ cells in RIPA lysis buffer supplemented with a protease/phosphatase inhibitor cocktail (HaltProtease and Phosphatase Inhibitor Cocktail). Protein concentration was determined using the bicinchoninic acid (BCA) protein assay per the manufacturer's protocol (ThermoFisher Rapid Gold BCA Protein Assay Kit). Equal amounts of protein (30μg) were resolved by Novex NuPAGE SDS-PAGE and transferred to low-fluorescent polyvinylidene fluoride (PVDF) membranes using a semi-dry transfer method (Invitrogen PowerBlotter). The membranes were then probed with antibodies against PSAT1 (1:1,000), PHGDH (1:1,000), ATF4 (1:500), GAPDH (1:2,000), COX-IV (1:5,000), and MGST1(1:500). A secondary goat anti-rabbit (1:15,000) was used. Immunoreactive bands were visualized using ECL or ECL+ reagent. WB densitometry was performed as describe in the literature (3).

**DNA constructs:** PSAT1 lentiviral constructs (Neo-EF1A-hPSAT1-T2A-TurboGFP) were designed to express Neomycin resistance, TurboGFP, and either the wild-type (WT) or enzyme-dead, structurally intact K200A (mut) PSAT1 protein. For PHGDH, constructs (Bsd-EF1A-hPHGDH-IRES-EBFP) expressed Blasticidin resistance, EBFP, and either WT or enzyme-dead D175N/R236K/H283A (mut) PHGDH protein. Lentiviral construct for overexpression of MGST1 was synthesized by VectorBuilder to express Neomycin resistance and TurboGFP along with human MGST, and the plasmid structure was the following: Neo-EF1A-hMGST1-IRES-TurboGFP. Constructs were synthesized by VectorBuilder. For PSAT1 knockouts, 3 Lentiviral constructs were synthesized by VectorBuilder carrying scrambled and two gene-targeted guide RNAs (gRNA30 and gRNA_exon1)(4) in the following structure: U6-gRNA-EFS-hCAS9-T2A-Puro.

**Mass spectrometry quantification of serine synthesis:** For HNT34 double rescued PSAT1 and PHGDH cells, 3 x $10^6$ cells were seeded in flasks and cultured under normal conditions for 2 days. On day 4, the media was replaced with fresh media supplemented with $^{13}$C-glucose. Cells were collected and centrifuged at 500 g for 5 mins at 4 degrees Celsius. The media was carefully removed, and the cell pellets were washed three times with 1 mL of PBS before being transferred to 1.5 mL safe-lock Eppendorf tubes. The cell pellets were centrifuged again at 500 g for 5 mins at

4 °C and the PBS supernatant was gently removed. Tubes then were snap-frozen in liquid nitrogen for 1 min and stored at -80°C. For controls, unlabeled cells for each cell clone, as well as parental HNT34 cells, were included. The same sample processing was performed for auxotrophic and non-auxotrophic cell lines (MUTZ3, YCUAML1 P31FUJ, UCSDAML1, MOLM13, OCIAML3, OCIAML5). Polar metabolites were extracted from samples with 80% methanol as the extraction solution. Polar metabolite-containing supernatant was isolated after centrifugation and dried under nitrogen gas. Dried metabolite extracts were resuspended in 50% methanol and then clarified by centrifugation at 15,000 g and 4 °C for 15 min. Finally, the supernatant was transferred into the 250ul sample vial and injected into the liquid mass spectrometry (LC-MS) system for analysis. LC–MS based metabolomics profiling was performed on a Vanquish UHPLC system (Thermo Fisher, Waltham, MA, USA) coupled with an Orbitrap Q Exactive™ HF-X mass spectrometer (Thermo Fisher, Waltham, MA, USA) at Complete Omics (Baltimore, MD, USA). The samples were injected into a Waters XBridge BEH Amide column (2.1 x 150 mm, 1.7μm) using a 25-min gradient at a flow rate of 0.2 mL/min. The LC parameters were as follows: autosampler temperature, 4 °C; injection volume, 2 ml; column temperature, 40 °C; and flow rate, 0.20 ml/min. The solvents and optimized gradient conditions for LC were: Solvent A, Water with 0.1% formic acid; Solvent B, Acetonitrile with 0.1% formic acid; A non-linear gradient from 99% B to 45% B in 25 minutes with 5min of post-run time. The QE HF-X mass spectrometer was operated in the negative mode with the following optimized operation parameters: Sheath gas flow rate, 35 arb; Aux gas flow rate, 10 arb; Sweep gas flow rate, 2 arb; Spray voltage, 2.85 kV; Capillary temperature, 350 °C. Data were acquired with Xcalibur acquisition software and processed with Compound Discoverer 3.3 (CD ver. 3.3, Thermo Fisher). The analyte database used for metabolites annotation was developed in-house with retention times based on the LC method described as above. The stable isotope labeling data were processed by the 'StableIsotopeLabeling' workflow unit of the Compound Discoverer software.

***In vivo*:** Animal experiments were approved by the Johns Hopkins University Animal Care and Use Committee. Six- to eight- week-old female NOD-SCID IL2Rgnull (NSG) mice were used for all *in vivo* experiments except of the experiment depicted in Figure 5C in which 6-week-old female NOD-scid IL2Rgnull-3/GM/SF (NSGS) mice were used. For leukemia burden assessment, peripheral blood (PB) was collected via tail vein bleeding on day 14, and then mice were euthanized for Bone Marrow (BM) and spleen harvest. BM cells were isolated from the femurs. The spleen, PB, and BM cells were incubated in the lysing buffer for 15 min × 3, then passed through a strainer into flow cytometry tubes. Cells were stained using anti-mouse CD45 conjugated with PE/Cy7 and anti-human CD45 conjugated with APC. Live/dead staining was performed using PO-PRO-1. Data analysis was performed using FlowJo software by directly comparing single human $CD45^+$ and single mouse $CD45^+$ events. Venetoclax was given orally in a total volume of 200 µL using a 20G feeding tube. The Venetoclax formulation consisted of 10% DMSO, 30% PEG400, and 60% Phosal-PG50.

**Methylation-specific qPCR:** Genomic DNA was isolated from 1 million cells using a genomic DNA extraction kit from New England Biolabs and subjected to bisulfite conversion using the EZ DNA Methylation-Gold Kit from Zymo Research. Primer pairs for unmethylated (unmeth) and methylated (meth) DNA were designed and validated for the correct size and methylation status through gel electrophoresis and Sanger sequencing. The qPCR was run in triplicate and captured on a BIORAD CPX96. The protocol is as follows: Initial denaturation: 95°C for 3 minutes, amplification: Denaturation: 95°C for 10 sec, annealing: 58.9°C for 30 sec (primers bind to target sequences). The total number of cycles was 45. Melt Curve Analysis: Increase by 0.5°C every 5 seconds starting from 65°C to 95°C

**Transcriptome and methylation analysis of primary AML cohorts.** The expression of PSAT1 mRNA levels according to different genetic aberrations in AML was performed by comparing RPKM levels obtained from cBioportal (https://www.cbioportal.org) for the TCGA and BEAT-AML cohorts. For the Leucegene cohort, TPM values and genetics were obtained from MISTIC (http://mistic.iric.ca), the Leucegene site (https://leucegene.ca), and GEO series GSE67040. Significance was assessed with unpaired t-tests on Box-Cox-transformed RPKM or TPM levels from the two groups (lambda optimized for normality by Kolmogorov-Smirnov testing). PSAT1 methylation values for comparison with PSAT1 mRNA were obtained from cBioportal for TCGA and from Giacopelli et al for BEAT-AML (5).

#### Supplementary Table 1

| | |
|---|---|
| **PSAT1 KO guides** | |
| sgRNA30 ($KO_1$) | CTTTCACAGAAATGAGTCAC |
| sgRNA_exon1 ($KO_2$) | CTTACTGAGTGCGGCAGCT (4) |
| **PSAT1 methylation specific primers** | |
| Methylated PSAT1 Forward primer | 5′- GTAGGGTTTGCGATAGTACGG -3′ (ref. 6) |
| Methylated PSAT1 Reverse primer | 5′- GCTACGATAAAAATCTACAACCGAC -3′ (ref. 6) |
| Unmethylated PSAT1 Forward primer | 5′- TAATTAGTGTAGGGTTTGTGATAGTATG -3′ |
| Unmethylated PSAT1 Reverse primer | 5′- AAAAATCTACAACCAACCCCAA -3′ |

### Supplementary Table 2

| Item | Brand | Catalog Number |
|---|---|---|
| ***Culturing*** | | |
| Fetal Bovine Serum (FBS), Premium, heat-inactivated | Thermo Fisher Scientific | A5670801 |
| Fetal Bovine Serum, dialyzed, US origin | Thermo Fisher Scientific | 26400044 |
| Serine | Millipore Sigma | S4311 |
| Glycine | Millipore Sigma | G8790 |
| Dextrose | Millipore Sigma | D9434 |
| Penicillin – Streptomycin | Thermo Fisher Scientific | 15140122 |
| RPMI 1640 Medium | Thermo Fisher Scientific | 11875093 |
| RPMI-1640 Media without Glucose, Glycine and Serine. 500mL, Sterile | Teknova | R9660-02 |
| Human plasma-like medium (HPLM) | Thermo Fisher Scientific | ME22541L1 |
| Human IL-6 Recombinant Protein, PeproTech® | Prepro tech / thermo | 200-06 |
| Human IL-3 Recombinant Protein, PeproTech® | Thermo Fisher Scientific | 200-03 |
| Human TPO (Thrombopoietin) Recombinant Protein, PeproTech® | Thermo Fisher Scientific | 300-18 |
| Human SCF, Animal-Free Recombinant Protein, PeproTech® | Thermo Fisher Scientific | AF-300-07 |
| Human Flt-3 Ligand (FLT3L) Recombinant Protein, PeproTech® | Thermo Fisher Scientific | 300-19 |
| Recombinant Human GM-CSF | Thermo Fisher Scientific | 300-03 |
| UM729 | Stemcell Technologies | 72332 |
| StemRegenin 1 (Hydrochloride) | Stemcell Technologies | 72352 |
| Vi-CELL BLU Reagent Kit | Beckman Coulter | C06019 |
| ***Genetics*** | | |
| SYBR™ Green Universal Master Mix | Thermo Fisher Scientific | 4309155 |
| EZ DNA Methylation-Gold Kit | Zymo Research | D5005 |
| Monarch® Genomic DNA Purification Kit | New England Biolabs (NEB) | T3010L |
| ***Animal experiments*** | | |
| HBSS | Thermo Fisher Scientific | 14025092 |
| Baker Amino Acid 1/2" Pellet IRR (5CC7) 5kg | Animal Specialties And Provisions | 1812426 |
| Mod TestDiet 5CC7 w/ No added Serine or Glycine 1/2" Pellet IRR (5BJX) 5kg | Animal Specialties And Provisions | 1817070-203 |
| Venetoclax | Selleck Chemicals | S8048 |
| Dimethyl sulfoxide | Millipore Sigma | D2650 |
| PEG400 | Selleck Chemicals | S6705 |
| Phosal 50 PG | MedChem Express | HY-Y1903 |
| ACK Lysing Buffer | Quality Biological | 118-156-101 |
| ***Flow cytometry staining*** | | |
| PE/Cy7 anti-mouse CD45 [30-F11] | BioLegend | 103114 |
| APC Mouse Anti-Human CD45 | BD Bioscience | 555485 |
| PO-PRO™-1 Iodide (435/455), 1 mM Solution in DMSO | Thermo Fisher Scientific | P3581 |
| eBioscience™ Propidium Iodide | Thermo Fisher Scientific | BMS500PI |
| ***Western Blotting*** | | |
| Psat1 Polyclonal Antibody | Thermo Fisher Scientific | PA5-22124 |
| Anti-PHGDH Antibody | Atlas Antibodies | HPA021241 |
| ATF-4 (D4B8) Rabbit mAb | Cell Signaling Technology | 11815S |
| GAPDH (D16H11) XP® Rabbit mAb | Cell Signaling Technology | 5174S |
| MGST1 Recombinant Rabbit Monoclonal Antibody (6A0S5) | Thermo Fisher Scientific | MA5-42631 |
| COX IV (3E11) Rabbit mAb | Cell Signaling Technology | 4850 |
| RIPA Lysis and Extraction Buffer | Thermo Fisher Scientific | 89900 |

| | | |
|---|---|---|
| Halt™ Protease and Phosphatase Inhibitor Cocktails, Thermo Scientific, Halt Protease and Phosphatase Inhibitor Cocktail, (100 X) | Thermo Fisher Scientific | 78440 |
| Pierce™ Rapid Gold BCA Protein Assay Kit | Thermo Fisher Scientific | 1863381 |
| NuPAGE™ LDS Sample Buffer (4X) | Thermo Fisher Scientific | NP0007 |
| NuPAGE™ Sample Reducing Agent (10X) | Thermo Fisher Scientific | NP0009 |
| NuPAGE™ Bis-Tris Mini Protein Gels, 4–12%, 1.0–1.5 mm | Thermo Fisher Scientific | NP0336BOX |
| NuPAGE™ Bis-Tris Mini Protein Gels, 4–12%, 1.0–1.5 mm | Thermo Fisher Scientific | NP0321BOX |
| NuPAGE™ Bis-Tris Midi Protein Gels, 4 to 12%, 1.0 mm | Thermo Fisher Scientific | WG1402BOX |
| Power Blotter Select Transfer Stacks, PVDF, regular size | Thermo Fisher Scientific | PB5310 |
| Power Blotter 1-Step™ Transfer Buffer (5X) | Thermo Fisher Scientific | PB7100 |
| Invitrogen™ NuPAGE™ MOPS SDS Running Buffer (20X) | Fisher Scientific | NP0001 |
| SuperBlock™ Blocking Buffer | Thermo Fisher Scientific | 37536 |
| Signalfire™ ECL Reagent | Cell Signaling Technology | 6883P3 |
| Signalfire™ Plus ECL Reagent | Cell Signaling Technology | 12630S |
| Pierce CL-Xposure™ Film, Thermo Scientific, CL-XPosure™ Film, Packaging=100 Sheets, Dimensions=12.7 x 17.8 cm (5 x 7) | Thermo Fisher Scientific | 34090 |
| Anti-rabbit IgG (H+L) (DyLight 680 Conjugate) | Cell Signaling Technology | 5366P |
| ***Cell transduction and Selection*** | | |
| LentiBOOST® Research Grade Lentivirus Transduction Enhancer Solution | Mayflower Bioscience | SB-P-LV-101-03 |
| Puromycin Dihydrochloride | Thermo Fisher Scientific | A1113803 |
| Geneticin Selective Antibiotic (G418 Sulfate) | Thermo Fisher Scientific | 10131035 |
| Corning® 50 mg Blasticidin S HCl | Corning | 30-100-RB |
| ***Software*** | | |
| GraphPad | Dotmatics | https://www.graphpad.com |
| FlowJo | BD | https://www.flowjo.com/ |
| SnapGene | Dotmatics | https://www.snapgene.com |
| MethPrimer | Ractigen Therapeutics | https://www.methprimer.com |
| ApE | M. Wayne Davis/ University of Utah | https://jorgensen.biology.utah.edu/wayned/ape/ |
| Fiji | Johannes Schindelin and collaborators | https://fiji.sc |
| Compound Discoverer™ | Thermo Fisher Scientific | https://www.thermofisher.com/us/en/home/industrial/mass-spectrometry/liquid-chromatography-mass-spectrometry-lc-ms/lc-ms-software/multi-omics-data-analysis/compound-discoverer-software.html |
| Microsoft Excel | Microsoft | https://www.microsoft.com/en-us/microsoft-365/excel/ |
| R | R Core Team | https://www.r-project.org |

### Supplementary Figure 1

A

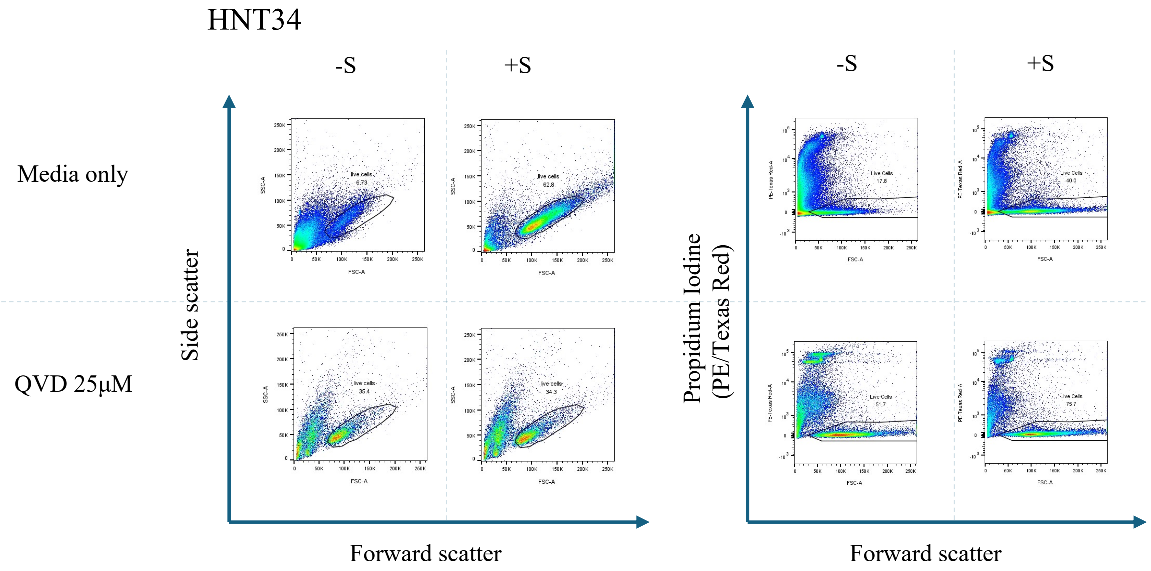


B

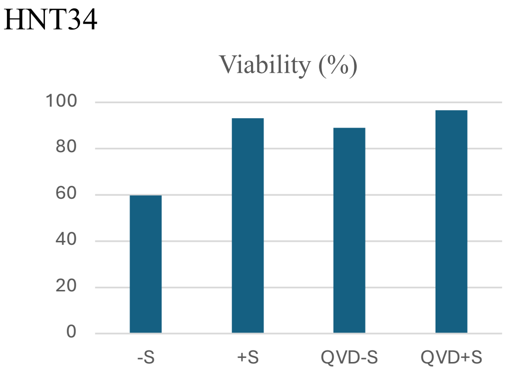


**Figure S1.** (A) Flow cytometry analysis of HNT34 cells after 3-day incubation in +S or –S RPMI, in the presence or absence of QVD (25 μM). The final DMSO concentration in QVD-treated conditions was 2.5%. Live cells were gated based on forward and side scatter or forward scatter and Propidium Iodine (PE/Texas Red channel) (B) Viability of HNT34 in the same conditions, assessed by trypan blue exclusion using the Vi-CELL BLU.

### Supplementary Figure 2

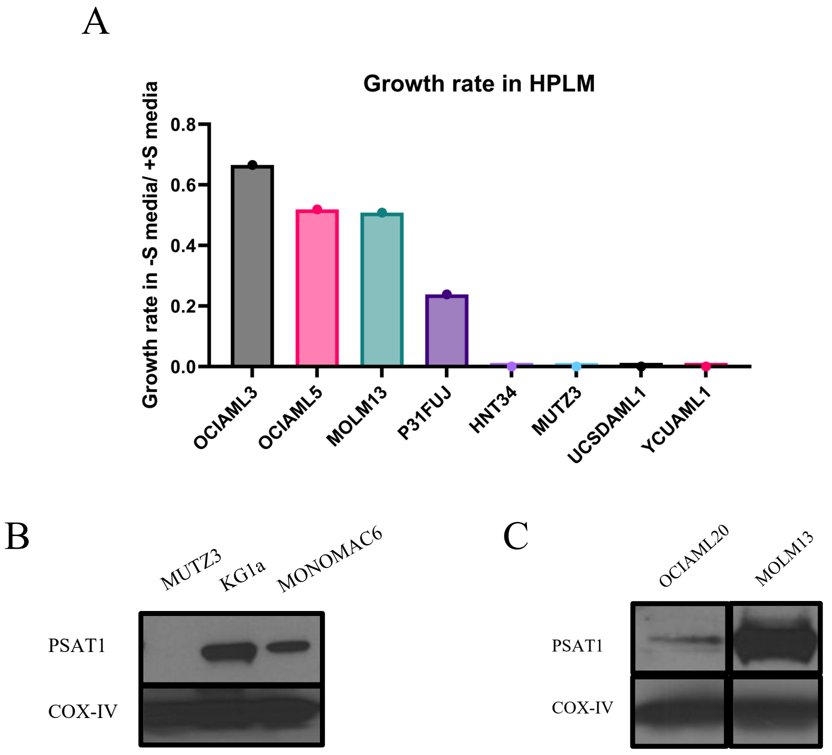


**Figure S2.** (A) AML growth rate (Malthusian constant; K) ratio as measured in starvation studies in -S and +S HPLM. Cell lines were run in triplicate or quadruplicate in +S and –S conditions for growth rate calculation. (B) Validated PSAT1 expression levels of MONOMAC6, using COX-IV as a loading control. KG1a serves as a positive and MUTZ3 as a negative PSAT1 control. (C) Increased sensitivity western blot showing limited PSAT1 expression in OCIAML20. MOLM13 was used as a PSAT1 positive control.

### Supplementary Figure 3

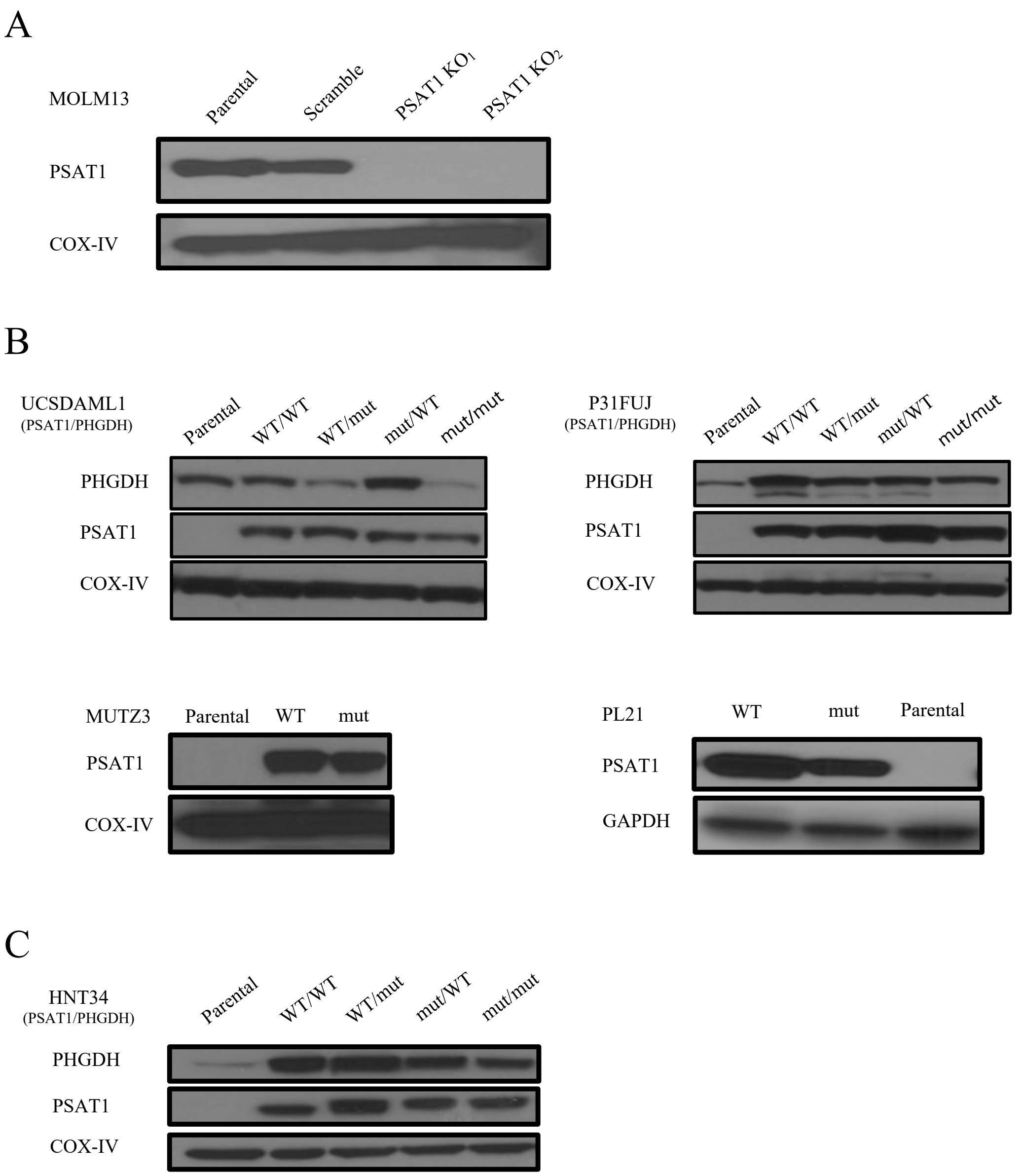


**Figure S3. Western blot validation** of (A) protein downregulation for PSAT1 in MOLM13 PSAT1 Knockout, and (B-C) upregulation of PSAT1 (mut or WT) in UCSDAML1, P31FUJ, MUTZ3, PL21, and HNT34, and PHGDH (mut or WT) in UCSDAML1, P31FUJ, and HNT34 transduced models. (B-C) "Mut" refers to inactivation mutations of the PSAT1 or PHGDH protein.

### Supplementary Figure 4

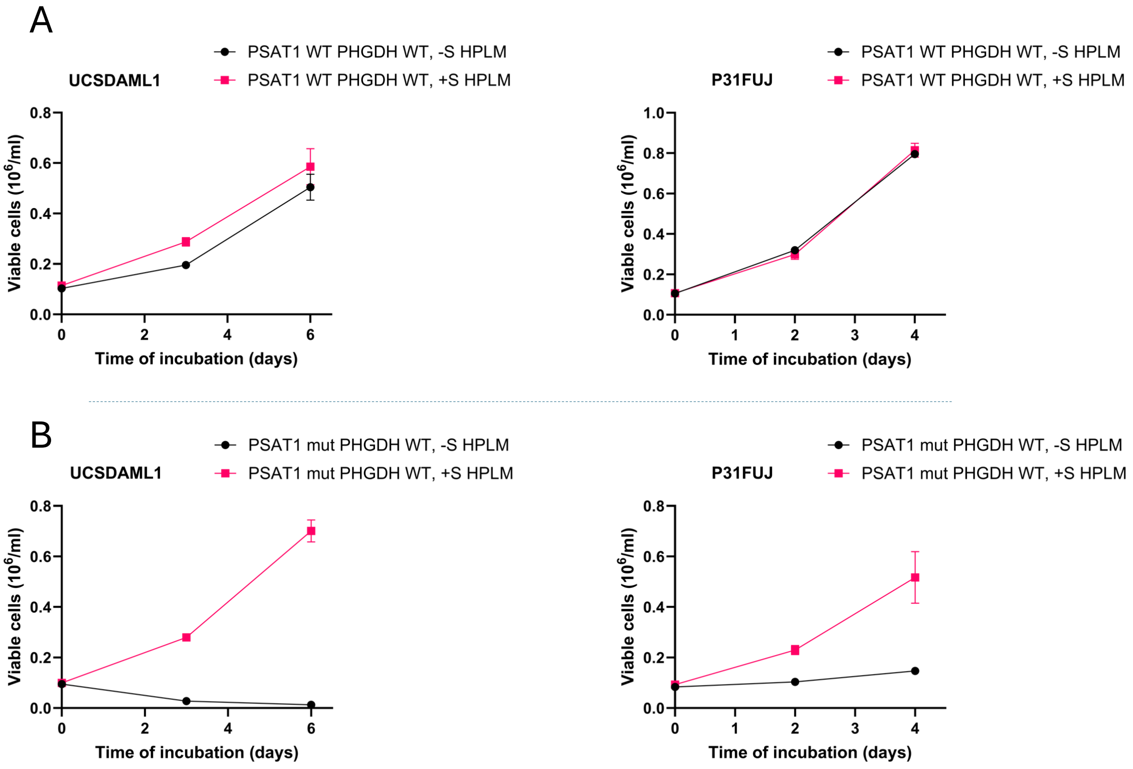


**Figure S4. PHGDH overexpression has no additive growth advantage in cells in serine-depleted conditions.** (A) Compared to the PSAT1 WT or mut rescue depicted in main Figure 4B, overexpression of PHGDH in UCSDAML1 and P31FUJ cells shows no additional growth benefit in –S HPLM. (B) PHGDH overexpression alone in UCSDAML1 and P31FUJ cells does not enhance growth in –S HPLM. Each sample was run in quadruplicate. “Mut” refers to inactivation mutations of the PSAT1 protein.

### Supplementary Figure 5

A

PSAT1 expression analysis

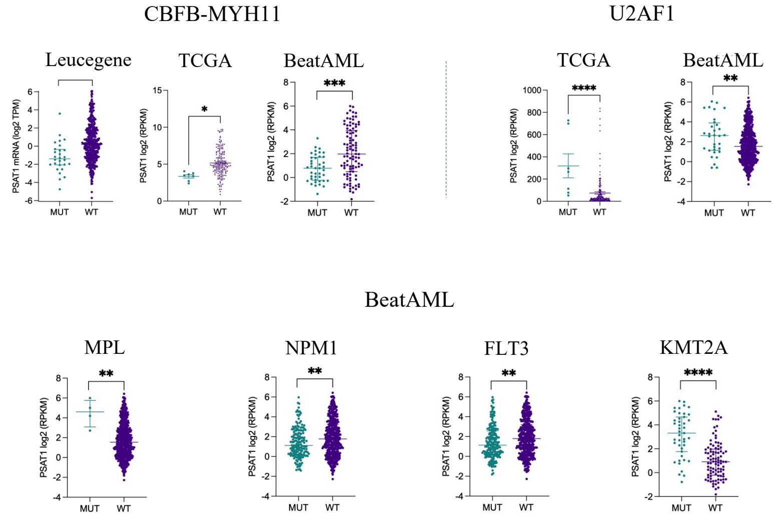


B

PSAT1 methylation analysis

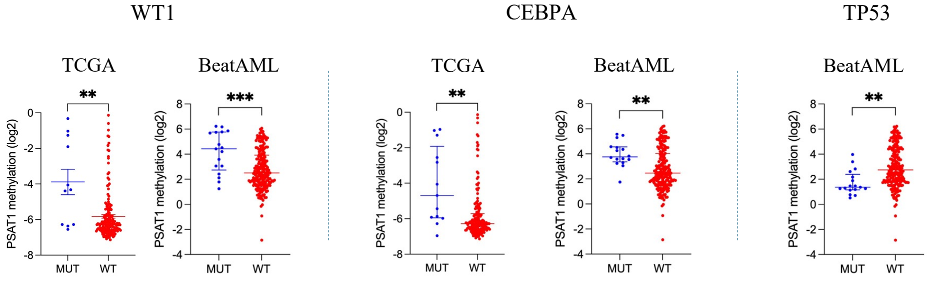


**Figure S5.** (A) PSAT1 expression analysis in AML patient samples harboring mutations in CBFB-MYH11, U2AF1, MPL, NPM1, FLT3, and KMT2A, using publicly available datasets from BeatAML, TCGA, and Leucegene. (B) PSAT1 promoter methylation analysis in AML patient samples with WT1, CEBPA, and TP53 mutations from the BeatAML and TCGA datasets.

### Supplementary Figure 6

A

p7314

+S -S

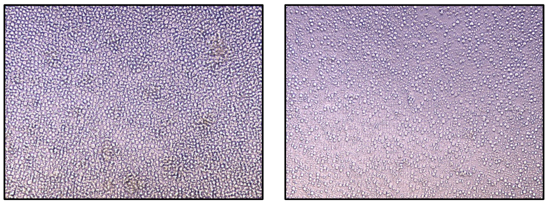

B

| | Cytogenetics | Mutations |
|---|---|---|
| p2112 | add(2)(q31),der(3)t(3;8)(q21;p21),der(8)t(2;8)(q?31;p21)[20] | PTPN11 p.G60V (9.81%)<br>SF3B1 p.K700E (41,71%) |
| p7314 | -7[16]/46,idem,+mar[4]<br>3q26(MECOM sep)*<br>-7(D7Z1,D7S486)x1* | - |
| p5548 | i(17)(q10[13]/47,sl,+mar[7] | KIT p.D816V (10.66%)<br>TP53 p.D281N (97.54%) |
| p4554 | normal | DNMT3A p.F752V (46.69%)<br>GATA2 p.A372T (32.02%)<br>NF1 p.S2414fs (23.54%)<br>GATA2 p.G200fs (11.66%) |
| p4319 | del(9)(q22q34)[20] | CEBPA p.Q312_K313insQ (or p.Q312dup) (29.05%)<br>WT1 p.S444fs (35.11%)<br>EZH2 p.Y244fs (36.48%) |
| p6965 | t(2;3)(p23;q27)[14] | SF3B1 p.K700E (31.04%) |
| p4229 | trisomy 8 (7% nuclei)* | WT1 c.1340-2A>G (splicing variant) (6.69%) |
| p4676 | normal | GATA2 p.A372T (32.06%)**<br>WT1 p.R441P (46.03%)**<br>WT1 p.S364fs (7.17%)** |
| p4359 | inv2(p21q13)[20] | FLT3 p.D835E (40.18%)<br>NPM1 p.W288fs (23.89%)<br>IDH2 p.R140Q (9.12%)<br>IDH1 p.R132C (35.31%)<br>DNMT3A p.R882H (46.15%) |

C

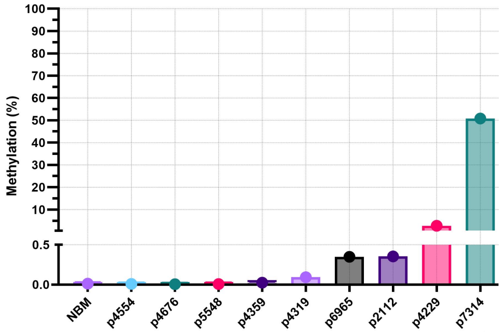

**Figure S6.** (A) Microscopy images showing a marked reduction in viability of the p7314 AML patient sample after 48h of incubation in +S and –S RPMI. (B) Cytogenetic (karyotype) and molecular data (NGS) from AML patient samples. The chromosomal translocation t(2,3)(p23;q27) in p6965 leads to an atypical MECOM rearrangement as described in the literature (7). *FISH findings, **NGS report obtained six months after initial sample collection. (C) Baseline DNA methylation levels (%) in AML patient samples, assessed by qPCR using the ΔΔCt method, comparing amplification from methylation-specific and unmethylation-specific primers. Each sample was analyzed in triplicate.
